# DLPFC Stimulation Suppresses High-Frequency Neural Activity in the Human sgACC

**DOI:** 10.1101/2025.03.26.645556

**Authors:** Ethan A. Solomon, Umair Hassan, Nicholas T. Trapp, Aaron D. Boes, Corey J. Keller

**Affiliations:** Dept. of Psychiatry and Behavioral Sciences, Stanford University Medical Center, Palo Alto CA 94305; Department of Neurology, Carver College of Medicine, University of Iowa, Iowa City IA 52242; Department of Psychiatry, Carver College of Medicine, University of Iowa, Iowa City IA 52242; Department of Pediatrics, Carver College of Medicine, University of Iowa, Iowa City IA 52242; Veterans Affairs Palo Alto Healthcare System, and the Sierra Pacific Mental Illness, Research, Education, and Clinical Center (MIRECC), Palo Alto CA 94305; Wu Tsai Neurosciences Institute, Stanford University, Stanford CA 94305

## Abstract

Transcranial magnetic stimulation (TMS) to the dorsolateral prefrontal cortex (DLPFC) is hypothesized to relieve symptoms of depression by inhibiting activity in the subgenual anterior cingulate cortex (sgACC). However, we have a limited understanding of how TMS influences neural activity in the sgACC, owing to its deep location within the brain. To better understand the mechanism of antidepressant response to TMS, we recruited two neurosurgical patients with indwelling electrodes and delivered TMS pulses to the DLPFC while simultaneously recording local field potentials from the sgACC. Spectral analysis revealed a decrease in high-frequency activity (HFA; 70-180 Hz) after each stimulation pulse, which was especially pronounced in the sgACC relative to other regions. TMS-evoked HFA power was generally anticorrelated between the DLPFC and sgACC, even while low-frequency phase locking between the two regions was enhanced. Together, these findings support the notion that TMS to the DLPFC can suppress neural firing in the sgACC, suggesting a possible mechanism by which this treatment regulates mood.

## Introduction

Transcranial magnetic stimulation (TMS) to the dorsolateral prefrontal cortex (DLPFC) is increasingly used as a treatment for major depressive disorder. Repetitive (rTMS) and theta burst stimulation (TBS) to the DLPFC have demonstrated clinically significant and durable improvement in depressive symptoms across clinical trials^1,2^, suggesting that this approach yields changes in the underlying brain networks that mediate mood disorders. Mechanistically, the prevailing hypothesis is that dysregulated activity of the subgenual anterior cingulate cortex (sgACC) contributes to depressive symptoms, as evidenced by reduction of sgACC activity in response to conventional treatments or through direct deep brain stimulation^3,4^. Though not accessible by TMS directly, modulation of sgACC activity via a node on the cortical surface – the DLPFC – could therefore be efficacious in depression. Specifically, investigators have targeted individualized subregions of the DLPFC with maximal functional MRI-based anticorrelation with the sgACC, under the assumption that putatively “excitatory” protocols to such a target would correspondingly inhibit activity of the sgACC^5^.

Clinical and fMRI evidence offers early support for this hypothesis. In retrospective analyses, patients with greater DLPFC-sgACC anticorrelation had better treatment outcomes^6–9^, and treatment itself can change the functional connectivity profile of regions linked to the sgACC^10^. However, further optimizing treatment protocols – and developing an enhanced mechanistic understanding of changes that occur with DLPFC TMS – requires electrophysiological evidence that goes beyond non-invasive neuroimaging. While fMRI BOLD can be a valuable measure, it is difficult to interpret when the hypothesized effects may involve the activity of inhibitory interneurons, which can both increase or decrease BOLD signal^11,12^.

Recent advances have allowed the application of TMS in neurosurgical patients with indwelling electrodes, allowing for the direct measurement of electrophysiologic activity from deep brain structures during stimulation. Using this approach, our group has previously reported that single pulse TMS to cortical targets provokes responses in a network of functionally-connected regions, including deep brain structures inaccessible to the direct effects of TMS^13,14^. One study examined the effects of TMS on the sgACC in the time domain, finding that DLPFC does cause evoked potentials in the sgACC^15^. However, neural responses in the sgACC may span the frequency spectrum, which is particularly important to investigate at higher frequencies if the hypothesized effect is inhibition of pyramidal neurons^16,17^. Here we closely examine the sgACC neural responses to single-pulse TMS (spTMS) to the DLPFC in two neurosurgical patients undergoing epilepsy neuromonitoring. We focus specifically on spectral phenomena, to offer an understanding of potential inhibitory effects of TMS that may not be apparent in analyses of evoked responses in the time domain alone.

## Results

### TMS-related change in sgACC spectral power

To characterize the electrophysiologic and spectral response in the sgACC to DLPFC TMS, we collected intracranial EEG (iEEG) from indwelling electrodes within sgACC in two neurosurgical patients; subject 460 underwent left-sided stimulation while subject 625 underwent right-sided stimulation (Figure 1A,E). TMS was delivered in a series of single pulses (50 pulses at 0.5 Hz) at 120% (subject 460) and 100% (subject 625) of the resting motor threshold, as well as a series of sham pulses (100 pulses at 0.5 Hz) in which the active side of the stimulator was rotated to face away from the scalp. Spectral methods were used to analyze the response in sgACC across frequency bands of interest (see Methods for details).

**Figure 1.**
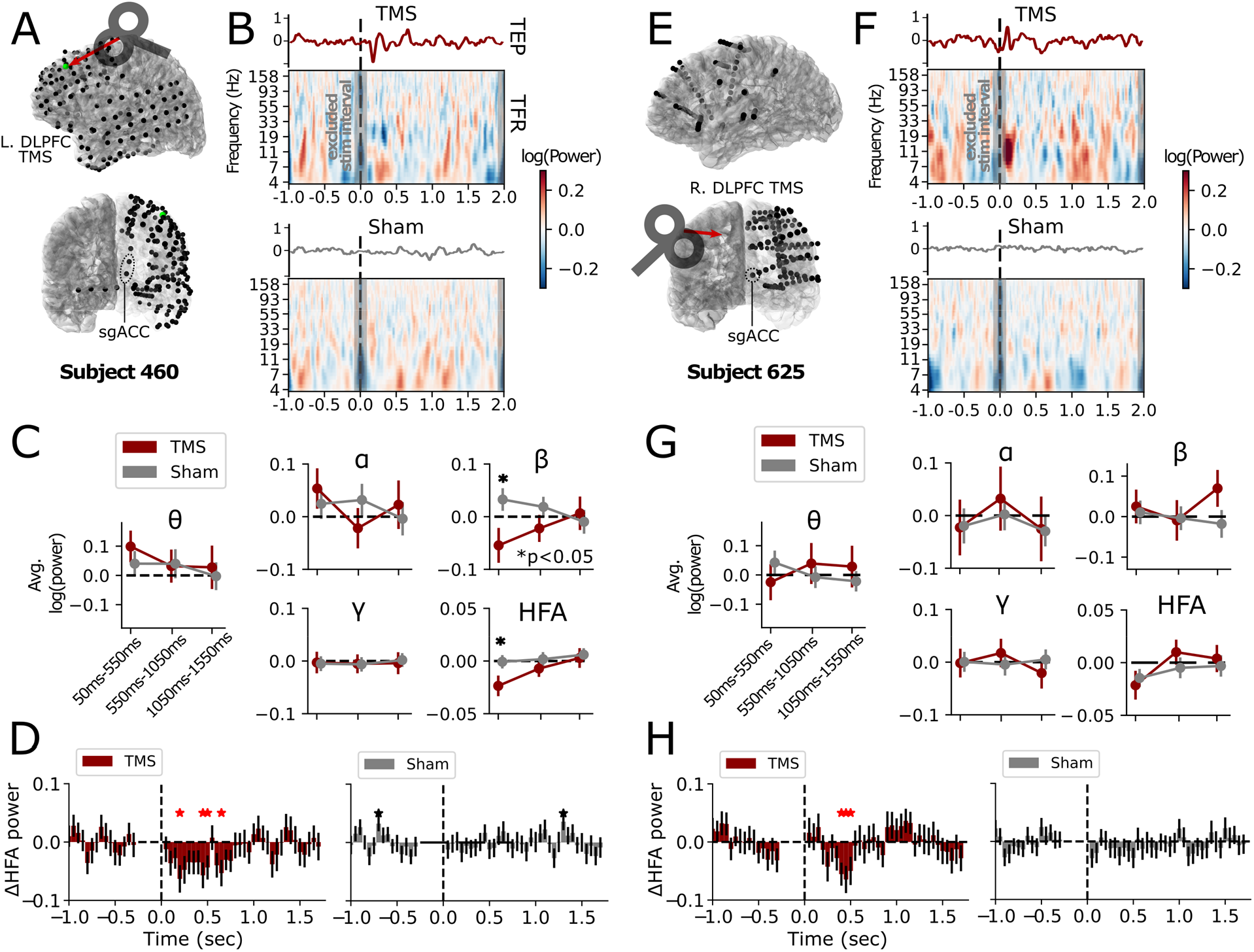
DLPFC TMS induces spectral power changes in the sgACC. **(A)** In subject 460, TMS was delivered to the left dorsolateral prefrontal cortex (DLPFC; green dot) in a series of single pulses spaced 2 seconds apart (0.5 Hz). Simultaneous intracranial EEG recordings were collected from subdural grids/strips as well as depth electrodes, including recording contacts in the left subgenual anterior cingulate (sgACC). **(B)** TMS-related activity in the sgACC was analyzed as both TMS-evoked potentials (top; TEP) and the spectral time-frequency response (bottom; TFR). Activity was analyzed separately for active TMS pulses and sham (active side of the coil pointed away from the skull; see Methods). Power values were normalized against the pre-stimulation (−1000 ms to -50 ms) baseline. **(C)** Spectral power was assessed across three 500 ms windows spanning 50-1550 ms post-stimulation, in key frequency bands of interest. In the first window (50-550 ms), spectral power during TMS trials was significantly lower than sham-related power (\**p*<0.05, uncorrected) in the beta (15-25 Hz) and HFA (70-180 Hz) ranges. **(D)** A time-resolved analysis was conducted for HFA power, examining effects within overlapping 200ms windows spaced 50ms apart. Significant TMS-related decreases in HFA were noted relative to pre-stimulation baseline in the approximate 0-750 ms interval (left). No similar effects were noted during sham trials (right). **(E-H)**: Same as A-D, for subject 625 who underwent right DLPFC stimulation and left sgACC recording. While no significant difference in power was found in any band at a coarse level of analysis (G), TMS-related HFA was significantly reduced relative to the pre-stimulation baseline in the 350-500 ms interval (H). Error bars show +/- 1 SEM.

In both subjects, single-pulse TMS to the DLPFC provoked detectable sgACC evoked potentials while sham did not, as has been previously reported (Figure 1B,F)^15^. In the frequency domain, active TMS decreased high-frequency activity (HFA; 70-180 Hz) spectral power relative to the pre-stimulation baseline in both subjects in the 50ms to 550ms post-stimulation interval (significant in subject 460: *t*(49)=-2.41, *p*=0.02; nonsignificant in subject 625: *t*(49)=-1.57, *p*=0.12). HFA suppression was significantly different from sham in subject 460 (2-sample *t-*test, *t*(148)=-2.013, p=0.046; Fig. 1C,G). In subject 460, a significant decrease in beta power (15-25 Hz) was also detected, relative to sham (*t*(148)=-2.27, *p*=0.024). These sham-corrected HFA and beta effects in subject 460 were not significant when FDR corrected for multiple comparisons over timepoints (HFA: corrected *p*=0.14; beta: corrected *p*=0.07). No other frequency bands demonstrated a significant TMS-related change in power (Fig. 1C,G).

To further examine dynamics of HFA suppression following DLPFC TMS, we extracted a measure of continuous HFA power from sgACC recording contacts. In subject 460, HFA was significantly suppressed relative to pre-stimulation baseline in the approximate 0-750 ms post-stimulation interval in the deepest sgACC contact (*p*<0.05, 1-sample *t*-test; Fig. 1D); similar measurements at non-sgACC contacts in more superficial locations along the same electrode shaft demonstrated shorter, fewer, or no significant intervals of HFA suppression (Fig. 2A).

**Figure 2.**
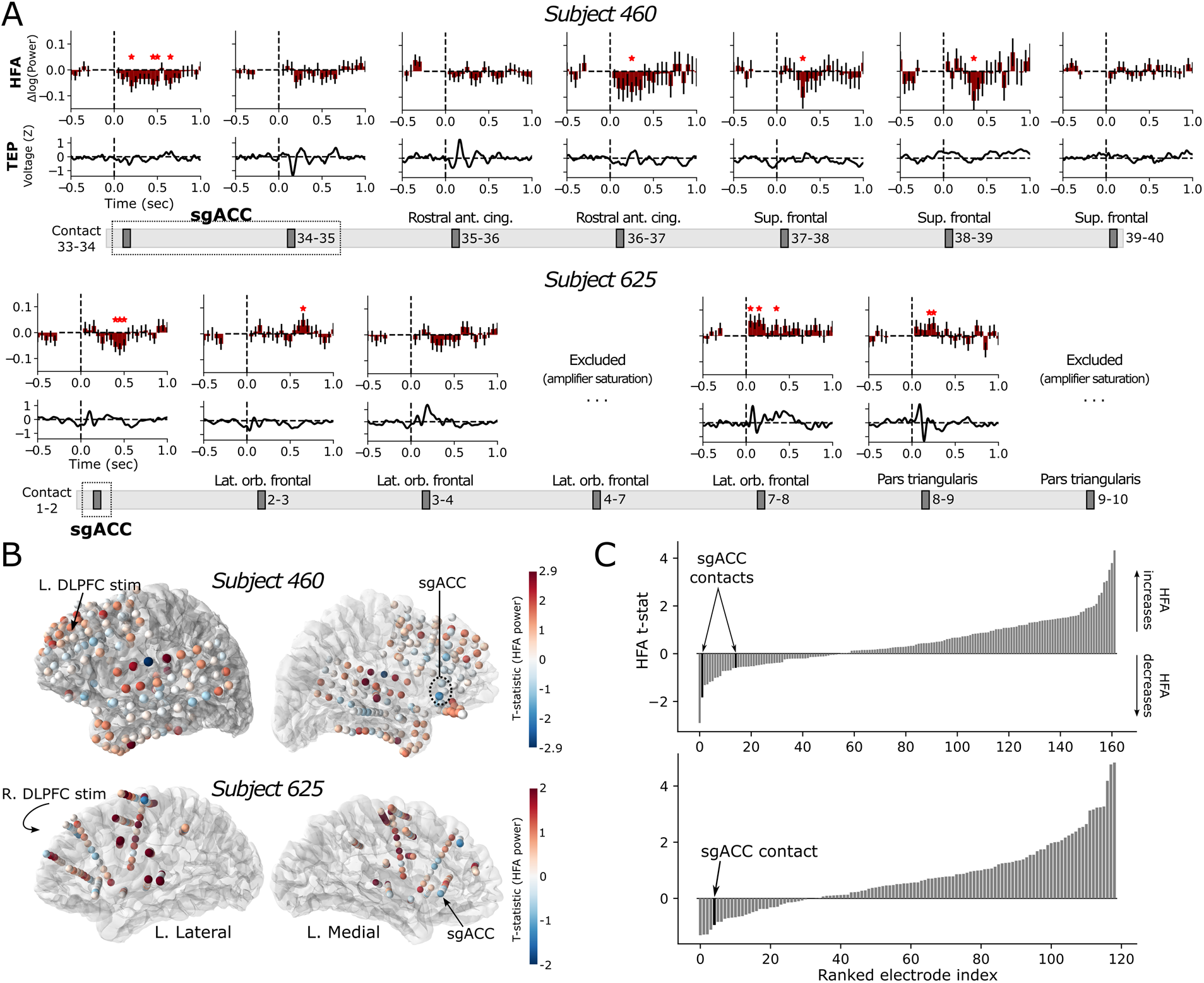
HFA suppression is pronounced in the sgACC relative to other brain regions. **(A)** Spectral power in the HFA range was extracted for each recording contact spanning the length of the depth electrodes that penetrated left sgACC (top: subject 460; bottom: subject 625). HFA was extracted in successive and overlapping 200ms windows, spaced by 50ms. While sgACC contacts in both subjects showed significant TMS-related decreases relative to baseline (\**p*<0.05 uncorrected, 1-sample *t-*test), the effect weakened or reversed at contacts closer to the cortical surface and further from sgACC. In subject 460, prominent TEPs were noted in contacts within (34-35) and adjacent to (35-36) the sgACC, while TEPs of varying morphologies were noted at all contacts spanning the depth electrode in subject 625. **(B)** A t-statistic was calculated reflecting the TMS-related change in HFA relative to baseline in the 50-550 ms interval, for each recording contact in each subject’s entire montage. Cooler colors correspond to greater suppression of HFA power relative to pre-stimulation baseline. **(C)** Ranked by degree of TMS-related change in HFA power, one sgACC contact saw the second-largest decrease in HFA power among all 162 recording contacts in the brain of subject 460 (top), and the fifth-largest decrease in HFA power in subject 625 (bottom).

Subject 625 showed qualitatively similar results and a significant (*p*<0.05, 1-sample *t*-test; Fig. 1H) reduction of HFA from baseline in the 350-500 ms interval, while more superficial contacts not located in sgACC demonstrated HFA increases (Fig 2A). HFA time-courses following sham stimulation showed no significant decrease from baseline (Fig. 1D,H)

Average HFA power was computed in the 50-550ms interval for all electrodes in the entire montage (Fig. 2B), revealing a distribution of HFA increases and decreases across the cortex and allowing comparison to sgACC effects. In subject 460, the deepest sgACC contact showed the second-largest magnitude HFA suppression of any contact in the entire brain, while sgACC in subject 625 showed the fifth-largest magnitude of HFA suppression (Fig. 2B,C).

The finding of HFA suppression in the sgACC raised the question as to how TMS affected DLPFC activity itself, and whether there was a link between HFA power in the two regions. We first focused on subject 460, who exhibited the strongest HFA modulation in sgACC and underwent left hemispheric stimulation and recording. HFA power time-courses were extracted from sgACC and DLPFC (by identifying a cortical surface contact near the stimulation coordinates). DLPFC and sgACC power time-courses showed an inverse relationship in post-stimulation power dynamics (Figure 3A-B; Pearson correlation, *r*=-0.60, *p*=0.0002). The same procedure was carried out between the sgACC and all other recording contacts in subject 460’s montage, generating a TMS-induced HFA power correlation *r* value for each contact. DLPFC contacts generally exhibited the strongest HFA anticorrelation with sgACC compared to other regions, though this effect was not universal across all DLPFC electrodes (Figure 3C-D, top). A similar effect was noted in subject 625, who underwent right-hemispheric TMS; specifically, contacts in the middle frontal gyrus demonstrated inverse HFA correlations with sgACC, while contacts in the superior frontal gyrus exhibited significant positive correlations (Fig. 3C-D, bottom). Following sham pulses, HFA was less robustly anticorrelated between DLPFC and sgACC compared to active TMS trials; maximal anticorrelation decreased to -0.43 (subject 460) and -0.32 (subject 625) across DLPFC recording contacts, and fewer DLPFC contacts overall exhibited anticorrelations with sgACC (Supplemental Figure 1). However, the presence of anticorrelated HFA sgACC-DLPFC activity was still notable in both participants (see Supplemental Figure 1C-D).

**Figure 3.**
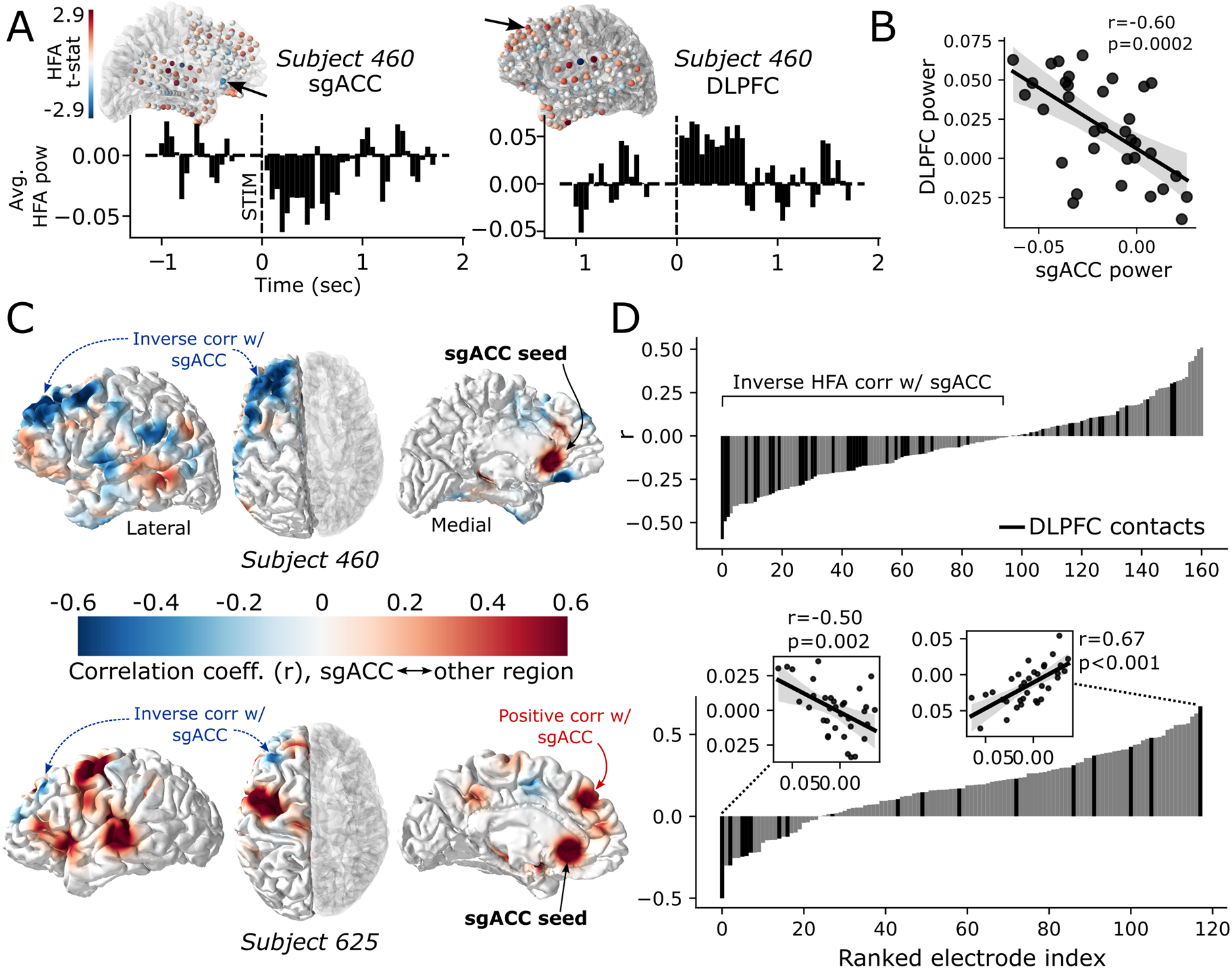
TMS-induced HFA power between sgACC and DLPFC is inversely correlated. **(A)** Baseline-corrected HFA power was averaged over trials to generate a time-course of activity in sgACC and DLPFC, as described previously (see Figure 2). HFA power values shown are for sgACC contact 33-34 and a DLPFC recording contact near the TMS stimulation site in subject 460. Contact locations are indicated on brain visualizations as in Figure 2B. **(B)** Across timepoints, HFA power values were inversely correlated between the sgACC and DLPFC contacts from panel (A) (Pearson correlation, *r*=-0.60, *p*=0.0002); shaded area shows 95% CI. **(C)** Using the sgACC contact as a seed, HFA power correlations were computed with all other contacts in the patient’s montage, and the resulting values projected to the cortical surface as a color map (see Methods for details). Blue values reflect inverse correlations with sgACC, while warm colors reflect positive correlations with sgACC. **(D)** All contacts in the montage were ranked according to HFA power correlations with the sgACC. Dark lines indicate recording contacts within the DLPFC, defined as the middle and superior frontal gyri in the Desikan-Killiany-Tourville (DKT) atlas. Insets show HFA sgACC power correlations of the maximally negative and positive DLPFC contacts.

Taken together, spectral power analysis revealed a TMS-related decrease in HFA power in both subjects, which appeared to be uniquely prominent in the sgACC relative to other brain regions. These results broadly align with prior findings from our group that TMS pulses can provoke fronto-temporal decreases in gamma and HFA power^13^, while confirming that the sgACC also exhibits a unique response profile beyond the more generalized spectral dynamics that have previously been reported. Moreover, TMS-provoked HFA in the sgACC was generally anticorrelated with frontal HFA activity, especially in more lateralized areas, and the DLPFC appeared to be a locus of this relationship relative to other regions in the brain.

### TMS-related change in sgACC spectral connectivity

While the inverse relationship in HFA between DLPFC and sgACC suggests these regions exhibit opposing patterns of neural activity, this finding alone does not explain how these regions communicate to achieve this antagonistic relationship. To uncover potential mechanisms of inter-regional communication, we examined whether DLPFC TMS shifted patterns of spectral connectivity between the targeted site and the sgACC, by analyzing the phase-locking value (PLV) between DLPFC contacts and sgACC contacts in the post-stimulation interval.

PLV reflects the degree to which there is a consistent *phase* relationship between two regions across all trials^18^. Of note, as no right-sided recordings were collected from subject 625, we analyzed PLV between the sgACC and two contacts of interest in this participant: (1) a left superior frontal gyrus contact as the closest frontal lobe recording site to the actual right-hemispheric TMS target (Figure 4), and (2) a recording contact in the homologous location in the left hemisphere corresponding to the right-sided target (Supplemental Figure 2).

**Figure 4.**
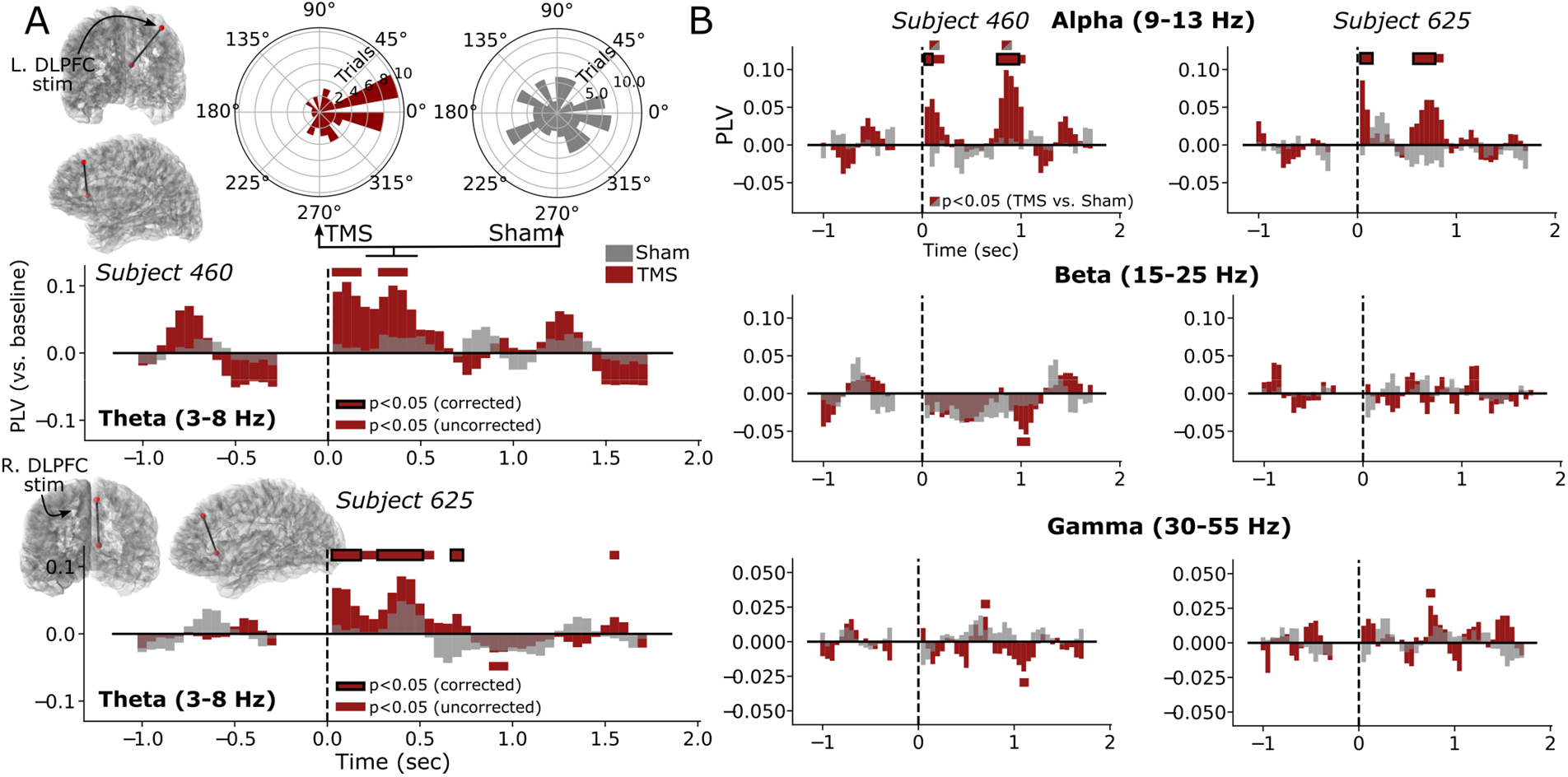
TMS-related change in spectral connectivity. **(A)** Connectivity between the DLPFC and sgACC was analyzed using the phase-locking value, a measure of phase consistency across trials. Changes in PLV were assessed in successive and overlapping 200ms windows, spaced 50ms apart, as done in power analyses. PLV values were statistically compared between TMS (red) and sham (gray) using a shuffling procedure (see Methods) and between pre- and post-stimulation intervals by z-scoring values against the pre-stimulation baseline period. Significant change (*p*<0.05) relative to baseline is indicated with red bars; a black border indicates significance after FDR correction. Red/gray boxes indicate time points of significant difference between TMS and sham. In the theta band (3-8 Hz), both subjects had significant PLV elevation above baseline in the approximate 50ms-500ms interval, but these differences did not significantly exceed sham. Polar distribution plots show the observed sgACC-DLPFC phase differences in subject 460 during an interval of significantly elevated theta PLV. **(B)** Across all frequency bands, alpha (9-13 Hz) showed prominent increases in sgACC-DLPFC phase locking in both participants, which were significantly different from sham in subject 460 (*p*<0.05, nonparametric; timepoints 100 ms and 850 ms). Briefer changes in beta and gamma band PLV did not survive FDR correction in either subject.

In both participants, we observed increased DLPFC-sgACC phase locking in the theta and alpha bands within the 1-second post stimulation period (Figure 4), but these increases were only significantly different from sham in the alpha band in subject 460 (timepoints 100 ms, 850 ms). Theta PLV increases above baseline were most prominent in the first 500 ms following stimulation, while alpha values demonstrated peaks in the first 100 ms and again from 750-1000 ms post-stimulation. These periods of enhanced connectivity approximately co-occurred with increased inter-trial phase coherence (ITC) within each electrode (Supplemental Figure 3), suggesting stimulation-related phase reset underlying increased inter-regional phase locking. PLV changes from baseline in the beta and gamma bands were brief and potentially driven by noise, as these effects did not survive multiple comparisons correction in either subject. In subject 625, significant theta and alpha DLPFC-sgACC phase locking (*p*<0.05, FDR corrected) was also observed using a contact in the homologous left frontal location corresponding to the right-sided TMS target, in the 50-150ms interval (Supplemental Figure 2).

## Discussion

Interactions between the DLPFC and sgACC are theorized to be essential to the antidepressant effect of TMS, and this hypothesis is backed by retrospective evidence from non-invasive neuroimaging^3,6–9^. However, the neural changes evoked by TMS have not been studied at the level of local field potentials, though such mechanisms may be important for understanding and optimizing the antidepressant effect of newer technologies. Leveraging intracranial neural recordings from two neurosurgical patients, we examined the sgACC neural response to single pulses of TMS to the DLPFC, finding that sgACC exhibited a marked decrease in HFA power following each pulse that was not evident to the same degree in other brain regions. Our results expand our understanding of how DLPFC TMS may exert its therapeutic effects.

We observed dynamic changes in the interactions between the DLPFC and the sgACC in response to DLPFC TMS, including enhanced low-frequency phase coupling following TMS pulses (Figure 4). During this period of enhanced coupling, the sgACC exhibited a marked decrease in HFA power. HFA is believed to be a general marker of aggregate cortical multiunit activity^16,17^, suggesting that the HFA decreases we measured in the sgACC reflect a suppression of population-level neural activity. This effect was observed to a greater degree in the sgACC relative to other brain regions. Moreover, post-stimulation HFA activity was generally anticorrelated between DLPFC and sgACC (Figure 3), such that greater HFA in the DLPFC was associated with suppressed HFA in the sgACC.

If the sgACC is a critical node in the network of brain regions regulating mood – and aberrant sgACC activation is correlated with depression – our results point to a possible mechanism by which TMS improves symptoms: activation of the DLPFC may in fact modulate the dynamic interactions between the DLPFC and sgACC, effectively enhancing DLPFC-associated inhibition of sgACC activity. Our finding that post-stimulation DLPFC HFA is inversely correlated with sgACC HFA also aligns well with existing hypotheses – and fMRI evidence – that DLPFC activity should be leveraged to exert an inhibitory influence on downstream regions like the sgACC^6,19^. The spectral evidence here hints that TMS provokes enhanced low-frequency phase coupling between the DLPFC and sgACC, which may be a transient mechanism by which the DLPFC causes an inhibitory effect on spiking activity.

However, our results also point to nuance in this story. The observed inverse correlation between DLPFC and sgACC HFA was notable, but far from universal – many DLPFC contacts exhibited positive power correlations, especially closer to midline frontal areas (though these could be rightly considered dorsomedial prefrontal cortex). These findings raise the possibility that targeting either positively or negatively correlated frontal regions could both yield therapeutic benefit, as has also been hypothesized^20^.

Our detailed spectral analysis adds valuable additional context to prior investigations of TMS-evoked phenomena in local field potentials. As shown here, observing a TMS-evoked potential is not necessarily a sign of “activation,” as these time-domain signatures can actually be associated with enhancement or suppression of activity in higher frequency bands (a useful proxy for population-level neural activity)^21^. Additionally, our observation of TMS-provoked phase locking in the theta/alpha bands deserves an important caveat: phase locking in these bands almost certainly reflects spectral overlap with the evoked potentials present in both the target area and downstream regions. As such, elevated PLV does not strictly mean TMS provokes an increase in phase-locked induced oscillations in both regions, but more likely suggests the low-frequency components of the TEP are closely aligned in time across these two regions.

Finally, our data, statistics, and conclusions are limited by our N=2 sample size. This sample is not sufficient to draw any strong or generalizable conclusions about TMS effects on the brain and inherently limits the robustness of our statistical approach. However, the consistent effects we observed across both subjects sets up intriguing questions for future research. Are there individuals where DLPFC TMS fails to alter sgACC activity, or even elevates it? Does the effect of TMS depend on the underlying brain state, similar to other closed-loop paradigms^22^? How would these effects differ if observed under TMS treatment protocols such as iTBS? The work here, while not yet generalizable, is a rare, valuable, and novel look into the mechanistic neural underpinnings of one of the most important new advances in depression treatment.

## Materials and Methods

### Human subjects

Two neurosurgical patients with medically intractable epilepsy underwent a surgical procedure to implant intracranial recording contacts on the cortical surface (electrocorticography) and within brain parenchyma (stereo-EEG). Contacts were placed in accordance with clinical need to localize epileptic regions. Each patient was admitted to the University of Iowa Hospitals and Clinics for up to 14 days of clinical and electrophysiological monitoring to identify their seizure focus. TMS experiments were conducted after the final surgical treatment plan was agreed upon between the clinical team and the patient, typically 1-2 days before the planned electrode explantation operation and 24 hours after the patient had restarted anti-seizure medications. All experimental procedures were approved by the University of Iowa Institutional Review Board, who reviewed safety data from a separate experiment prior to approval for human subjects. Written informed consent was obtained from both participants.

### Imaging protocol and intracranial electrode localization

Intracranial electrodes were localized in a manner identical to that previously described in Wang, et al.^14^ Patients underwent anatomical and functional MRI scans within two weeks of electrode implantation. The day following implantation, subjects underwent a second MRI and volumetric computerized tomography (CT) scans. The location of each contact was identified on the post-implantation T1-weighted MRI and CT, and subsequently post-implantation scans were transformed to pre-implantation T1 anatomical space in a manner that accounts for the possibility of post-surgical brain shift. Freesurfer^23^ was used to map electrode locations onto a standardized set of coordinates across subjects, which were then labeled according to their location within the Desikan-Killiany-Tourville (DKT) anatomical atlas^24^. The location of sgACC contacts was manually confirmed on postoperative imaging by an expert neuroanatomist (A. Boes).

### Transcranial magnetic stimulation

For stimulation, we used a MagVenture MagVita X100 230V system with a figure-of-eight liquid-cooled Cool-B65 A/P coil (Magventure; Alpharetta, GA, USA). Stimulation pulses were biphasic sinusoidals with a pulse width of 290 microseconds, with stimulator output set at a percentage of each subject’s motor threshold. Pulses were delivered at 0.5Hz, allowing for 2-second inter-stimulation intervals to examine spectral responses. TMS experiments were conducted 12-13 days after implantation and after starting anti-seizure medications. Neuronavigation using frameless stereotaxy was guided with Brainsight software supplied with the pre-implantation T1/MPRAGE anatomical scan. Stimulation parameters were recorded in Brainsight during all experimental trials. Motor thresholds were determined starting with the hand knob of the motor cortex as a target, beginning at 50% machine output and adjusted in small increments until movements were observed in 50% of trials.

Single TMS pulses were directed at DLPFC at 120% resting motor threshold in subject 460 and 100% resting motor threshold in subject 625. DLPFC targets were defined by the Beam F3 region^25^, identified by transforming published coordinates (MNI 1mm: -41.5, 41.1, 33.4) into each subject’s native T1 and displaying it in Brainsight; the corresponding location on the right hemisphere was used for subject 625. The stimulation site was adjusted slightly if access was impeded by head wrap or anchor bolts for securing electrodes. Sham pulses were delivered in an identical manner to active, with the TMS coil flipped 180 degrees such that the magnetic field was directed away from the head. Participants underwent 50 stimulation pulses (“trials”) and 100 sham pulses delivered at 0.5 Hz (i.e. 2-second inter-stimulation interval) automatically using the BEST toolbox^26^.

### iEEG recording

Electrode recordings were conducted in a manner identical to Wang, et al.^14^ and detailed in Hassan, et al.^27^. Briefly, depth and grid electrodes (Ad-Tech Medical; Racine, WI, USA) were either stereotactically implanted or placed on the cortical surface, respectively. A platinum-iridium strip electrode placed in the midline subgaleal space was used as a reference. Data were amplified, filtered (ATLAS, Neuralynx, Bozeman MT; 0.7-800 Hz bandpass), and digitized at 8000 Hz. In both subjects, contacts were excluded from analysis if stimulation artifact saturated the amplifier, if electrodes were placed within an identified seizure onset zone, or if electrodes were contaminated by non-neural noise indicative of poor connection or placement outside the brain.

### iEEG preprocessing and power analysis

The FieldTrip MATLAB toolbox^28^ was used to load iEEG data into our analysis pipeline. Data preprocessing and analysis was principally done with the MNE toolbox in Python^29^. Raw signals were initially re-referenced to account for large-scale noise or contamination of the reference electrodes; stereo-EEG (depth) electrodes were re-referenced using a bipolar montage, while grids and strips on the cortical surface were collectively re-referenced to their common average.

Although we avoided statistical analysis of the period of time containing the ∼15 ms stimulation artifact itself, the use of spectral methods continuous over time necessitates use of the full iEEG signal during each trial. For this reason, we scrubbed the stimulation artifact from all signals and replaced it with synthesized stationary iEEG that reflects a similar spectral profile as the background^13,30^. Specifically, the iEEG signal was clipped from 25 ms prior to 25 ms following stimulation and replaced with a weighted average of the 50 ms immediately following and prior to stimulation. The pre- and post-stimulation clips were first reversed, then tapered linearly to zero along the length of the signal, and then finally summed together to replace the artifact period. Finally, signals were notch filtered at 60 Hz and harmonics to remove line noise, using an F-test to find and remove sinusoidal components. Lastly, signals were downsampled to 500 Hz for further analysis.

To statistically compare spectral activity in TMS trials against sham trials – in order to control for auditory and expectancy effects associated with the stimulation click^31^ – iEEG signals from each contact in the dataset were first segmented into 3-second intervals, spanning 1 second prior to stimulation to 2 seconds after stimulation. To extract measures of spectral power, we used Welch’s method to measure the power spectral density (PSD) from 3 Hz to 180 Hz. Spectral decomposition was performed in either 500 ms windows (Figure 1D,H) or 200 ms windows (Figure 2) spanning the iEEG signal, depending on the granularity of analysis. 200 ms windows were spaced by 50 ms, providing some overlap between each successive window. Next, we log-transformed and then averaged power values into frequency bands of interest: theta (3-8 Hz), alpha (9-13 Hz), beta (15-25 Hz), gamma (30-55 Hz), and high frequency activity (HFA; 70-180 Hz). 200 ms windows were only used for HFA power analyses; all windows overlapping the stimulation pulse timepoint were excluded from further analyses.

Finally, for each trial, baseline power (averaged in the -1000 ms to -200 ms range, to avoid overlap with stimulation artifact) was subtracted from all subsequent power, correcting for natural drift in baseline power over time. Baseline-corrected powers were compared between TMS trials and sham trials using two sample *t*-tests, generating a *t*-statistic for each time window, frequency band, and recording contact in both subjects. Baseline-corrected power values were also statistically compared to zero (Figure 2) using 1-sample *t*-tests.

### Surface visualizations

Brain surface maps of TMS-iEEG effects (Figure 2B, 3C) were generated using the PySurfer library^32^. The value corresponding to each recording contact was smoothed using a Gaussian kernel (sigma=6) and then projected to each subject’s individual cortical surface (extracted via FreeSurfer, see *Imaging protocol and intracranial electrode localization* section above) with the overall effect of any individual electrode limited to a radius of 20. Projected values were colored according to their magnitude.

### Comparison of DLPFC and sgACC HFA power

To analyze the relationship between HFA power in the DLPFC and sgACC (Figure 3), trial-averaged HFA power was extracted in successive and partially overlapping 200 ms windows for all recording contacts in the brain, in the same manner as was done for sgACC power (Figures 1-2). The power values across all baseline-corrected and post-stimulation 200 ms windows were compared (Pearson correlation) between the sgACC and all other contacts in the brain. DLPFC contacts were defined as those in the superior frontal gyrus, rostral middle frontal gyrus, or caudal middle frontal gyrus in the Desikan-Killiany-Tourville atlas. The DLPFC contact with the maximal inverse correlation with sgACC in subject 460 is shown as an example in Figure 3A-B; note that this is the second-closest contact to the actual spTMS target site.

### Phase-locking value (PLV) connectivity analysis

The phase-locking value^18^ was analyzed by first using the Morlet wavelet method to generate a continuous measure of spectral phase across each trial, with evenly-spaced wavelets spanning each frequency band of interest. The number of cycles was adjusted for each band (theta, 2; alpha, 3; beta, 4; gamma, 5). HFA was avoided as a band of interest for connectivity given substantial prior evidence that long-range connectivity is not detectable in this range via standard macroelectrodes.^33,34^

PLV was computed as the concentration of phase differences between two recording contacts, measured across all trials, via the MNE implementation. Recording contacts were chosen as (1) the deepest sgACC contact and (2) the DLPFC (see anatomical definition above) contact closest to the actual spTMS target; note that in subject 625, who underwent right-hemispheric stimulation – this was an electrode in the left superior frontal gyrus closest to that right-sided target. (The homologous left-sided location was also analyzed and is presented in Supplemental Figure 2.) Baseline PLV was averaged in the -950 ms to -250 ms range and subtracted from subsequent values, to generate a continuous measure of TMS- or sham-related change in PLV (Figure 4). To assess for statistical deviation from baseline, PLV values were z-scored relative to values in the baseline interval, which was then used to get a corresponding *p*-value via the cumulative normal distribution.

To assess for statistical difference between TMS and sham-related PLV, a non-parametric shuffling procedure was used to account for the bias introduced by differing numbers of TMS and sham trials. TMS and sham trials were shuffled randomly and divided into groups of 50 and 100 trials, to serve as surrogate values in creating a null distribution. PLV was measured in each surrogate group, then subtracted from one another to get a measure of PLV-difference expected under chance. This procedure was repeated 100 times, generating a null distribution of chance-level PLV differences. Finally, the true PLV difference was compared to this distribution to generate a *p*-value. Further details on this procedure have been published elsewhere^35^.

The analysis of inter-trial phase coherence (ITC) presented in Supplemental Figure 3 was carried out in a similar manner to PLV, using the consistency of measured phase from each electrode individually (instead of phase differences between two electrodes).

## Acknowledgments

We thank Matt Howard and Ben Pace for their contributions to imaging acquisition and data collection efforts, and Charlie Dickey for valuable feedback on this manuscript. We also thank the neurosurgical patients who selflessly participated in this research.

## Author Contributions

*Ethan Solomon*: Conceptualization, formal analysis, software, data curation, writing – original draft, writing – review and editing, visualization. *Umair Hassan*: Formal analysis, data curation, writing – review and editing. *Nicholas Trapp*: Conceptualization, investigation, resources, data curation, writing – review and editing. *Aaron Boes*: Conceptualization, methodology, investigation, resources, supervision, project administration, funding acquisition, writing – original draft, writing – review and editing. *Corey Keller*: Conceptualization, methodology, resources, supervision, project administration, funding acquisition, writing – original draft, writing – review and editing.

