## Supplemental Figures for "DLPFC Stimulation Suppresses High-Frequency Neural Activity in the Human sgACC"

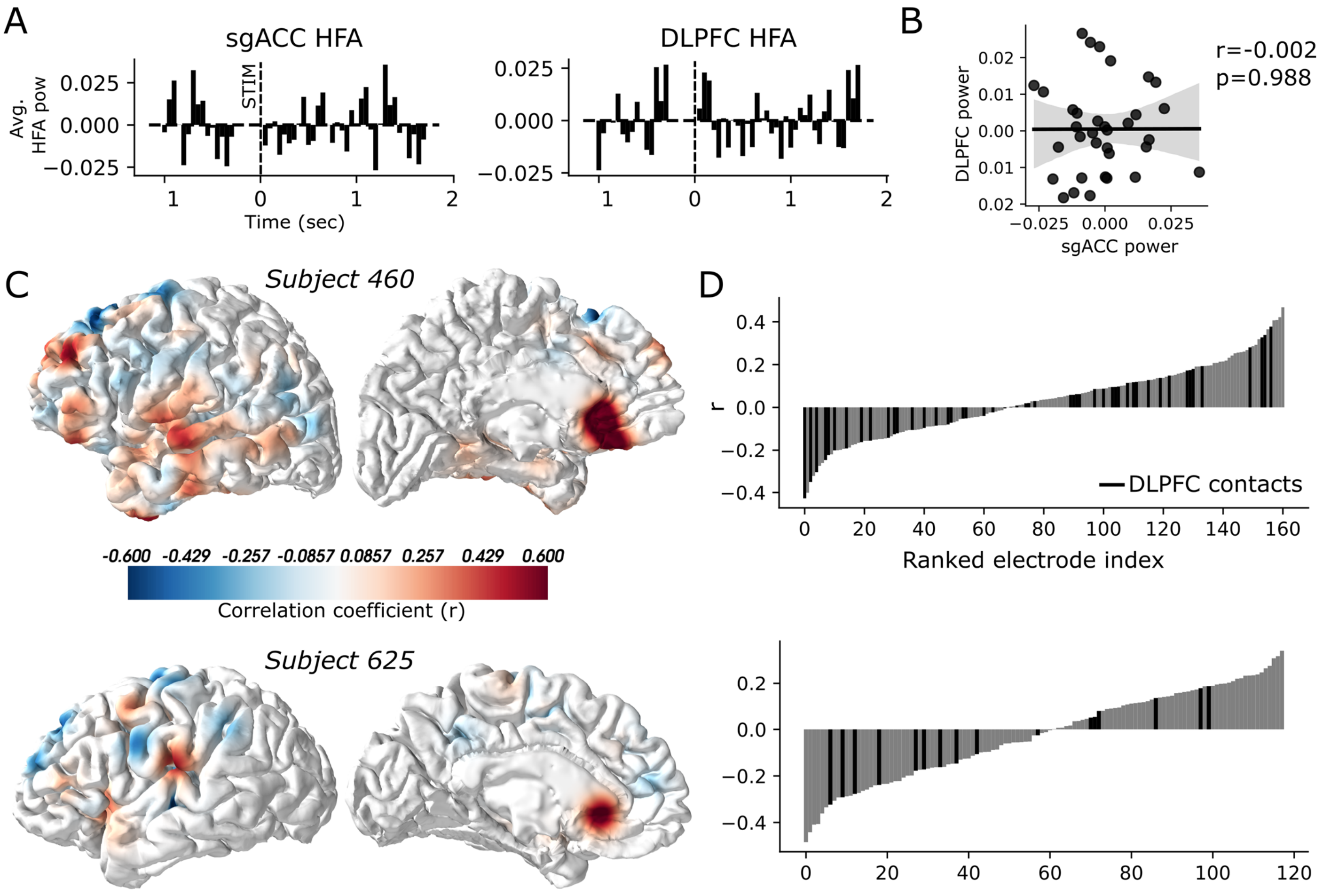


**Supplemental Figure 1. sgACC-DLPFC HFA anticorrelation is present, but less robust, following sham spTMS.** Figure structured identically to Figure 3 in the main text, analyzed only for sham trials. **(A)** HFA power time-courses following sham spTMS in the same recording contacts highlighted in Figure 3A. **(B)** In the same pair of DLPFC-sgACC contacts highlighted in Figure 3, there is no significant correlation between HFA power values (Pearson correlation, *r*=-0.002, p=0.988) during sham. **(C)** As in Figure 3C, the correlation *r* values between all recording contacts and the sgACC were projected as colors onto the cortical surface of both subjects. Blue areas indicate inverse sgACC-DLPFC correlations. **(D)** As in Figure 3D, all recording contacts were ranked according to correlation with the sgACC. DLPFC electrodes are indicated in black.

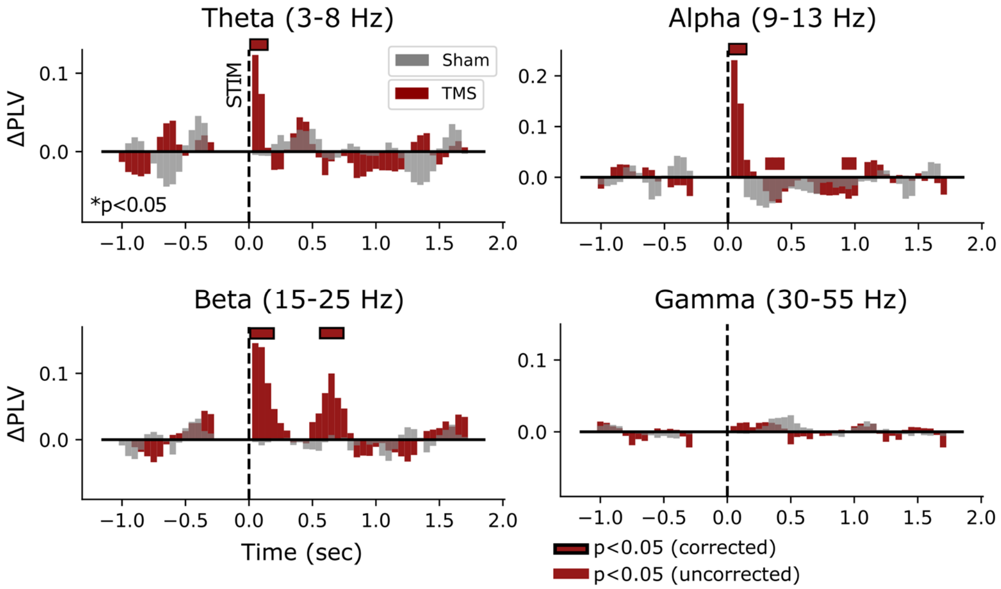


**Supplemental Figure 2. sgACC to DLPFC phase locking value in subject 625 on left-sided homolog of right-sided stimulation site.** Structured as in Figure 4A,B, using a left DLPFC contact in the homologous location to the right-sided TMS site. *p*<0.05 uncorrected (red), FDR corrected (red with black border).

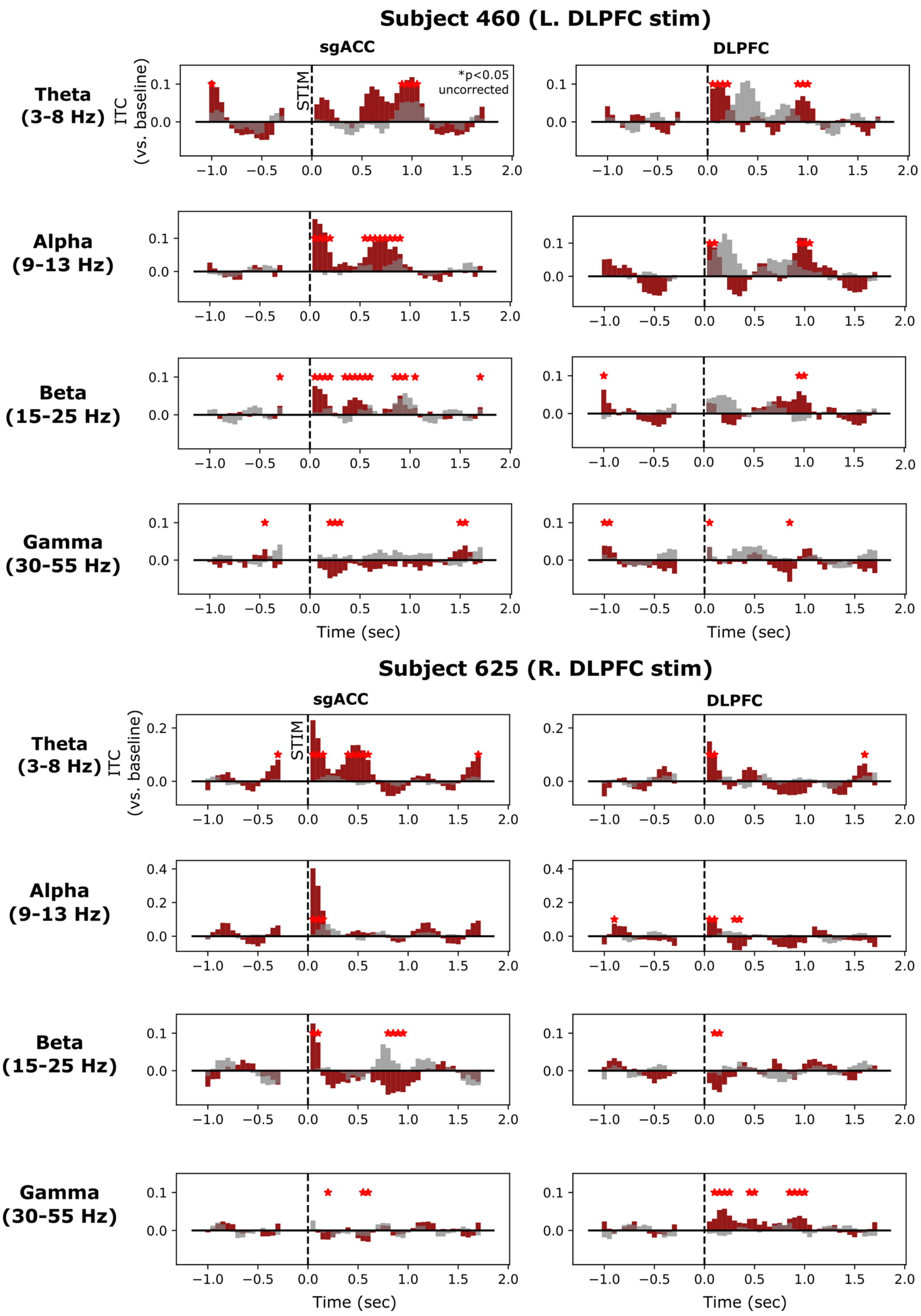


**Supplemental Figure 3. Inter-trial phase coherence (ITC) is modulated by single pulse TMS.** In subject 460 (top) and subject 625 (bottom), inter-trial phase coherence was calculated as the phase consistency of a given frequency, across all timepoints. Higher values reflect a similar phase from trial-to-trial. In both the sgACC and DLPFC, theta and alpha bands demonstrated the greatest change in ITC relative to pre-stimulation baseline, but notable increases were also observed in the beta band (subject 460 sgACC) and gamma band (subject 625 DLPFC). *$p$<0.05, uncorrected.
